# Diversity and prevalence of ANTAR RNAs across actinobacteria

**DOI:** 10.1101/2020.10.11.335034

**Authors:** Dolly Mehta, Arati Ramesh

## Abstract

Computational approaches are often used to predict regulatory RNAs in bacteria, but their success is limited to RNAs that are highly conserved across phyla, in sequence and structure. The ANTAR regulatory system consists of a family of RNAs (the ANTAR-target RNAs) that selectively recruit ANTAR proteins. This protein-RNA complex together regulates genes at the level of translation or transcriptional elongation. Despite the widespread distribution of ANTAR proteins in bacteria, their targets RNAs haven’t been identified in certain bacterial phyla such as actinobacteria. Here, by using a computational search model that is tuned to actinobacterial genomes, we comprehensively identify ANTAR-target RNAs in actinobacteria. These RNA motifs lie in select transcripts, often overlapping with the ribosome binding site or start codon, to regulate translation. Transcripts harboring ANTAR-target RNAs majorly encode proteins involved in the transport and metabolism of cellular metabolites like sugars, amino acids and ions; or encode transcription factors that in turn regulate diverse genes. In this report, we substantially diversify and expand the family of ANTAR RNAs across bacteria.

## INTRODUCTION

Actinobacteria is a ubiquitous bacterial phylum, widely distributed across terrestrial and aquatic ecosystems (Parks et al. 2018). The phylum consists of very diverse bacteria, ranging from defensive mutualists dwelling in varied habitats to gastrointestinal commensals that provide beneficial properties to their host. They are also the largest source of novel natural antibiotics, enzymes and secondary metabolites. In addition to their immense environmental and industrial impact, this phylum also consists of pathogens such as species from *Corynebacterium, Nocardia, Mycobacterium* and *Rhodococcus*, which cause disease in humans, animals and plants (Ventura et al. 2007).

The diversity of environmental niches seen within the actinobacteria phylum argues for diverse mechanisms of gene regulation that would allow an efficient response to environmental changes. While a body of literature now places non-coding RNAs and RNA-protein based mechanisms as a major mode of gene-regulation in several model bacteria, our knowledge of RNA-based regulatory mechanisms in actinobacteria remains limited.

One approach to identifying regulatory RNAs in actinobacteria, has been using deep sequencing of the transcriptome coupled with 5’-RACE mapping, to identify potential RNAs that map to the untranslated regions (UTRs). These RNAs are then subjected to structure prediction tools (Washietl et al. 2005; Vitreschak et al. 2001; Lorenz et al. 2011) and compared against known RNA families to confirm the presence of regulatory RNAs. This approach in *Corynebacterium and Streptomyces* under exponential growth conditions has led to the identification of 6C RNA family, 6S RNA family, T-box leader element and several known riboswitches such as yybP-ykoY, TPP, FMN, SAM and cobalamin, in addition to novel sRNAs and trans-encoded RNAs (Mentz et al. 2013; Vockenhuber et al. 2011). A similar approach led to the discovery of 75 novel small RNAs in *Rhodococcus sp*. when grown in glucose and pyrene as sole carbon sources, a small fraction of which have now been assigned functions (Peng et al. 2020). Such an approach requires cells to be grown under specific conditions of interest, and do not identify the repertoire of RNAs that the cell can produce in response to unknown signals and cues.

Computational methods have also been successfully employed, to identify regulatory RNAs in actinobacteria. In one study, homologs of genes were first identified and their upstream intergenic regions were aligned and searched for patterns/ motifs using RNA secondary structure prediction tools such as RNA-pattern (Vitreschak et al. 2001) and PAT (A.V.Seliverstov, unpublished). This led to the identification of LEU element (Seliverstov et al. 2005), T-box (Seliverstov et al. 2005) and B12 (Vitreschak et al. 2003) riboswitches in several actinobacteria. More generally, the RNA family database (Rfam) employs covariance analysis, wherein bacterial genome sequences are scanned for conserved base-pairing patterns, to identify structurally conserved RNA families in the genome. Based on this, the Rfam database suggests the presence of ∼90 cis-regulatory RNA families in one or more actinobacteria (Rfam v14.2). While these approaches have identified RNAs in actinobacteria, they are mostly limited to RNA families that are highly conserved in sequence and structure, where homologs from different bacterial phyla closely resemble each other.

For some RNA families, the highly GC rich actinobacterial genomes may result in RNA sequences that are diverged from their firmicute or proteobacterial homologs, and hence not easily identified through routine sequence based or structure based searches. One such example is the 6S RNA family, which could only be identified in actinobacteria using a clustering method wherein the sub-optimal RNA structures were used to find functionally relevant motifs (Pánek et al. 2011). Known 6S RNAs from related bacterial species of proteobacteria, firmicutes and cyanobacteria were analyzed for similarity based on sequence and minimum free energy (MFE) structures. Despite a common function, these RNAs lack sequence and structure similarity. Instead of MFE structure when sub-optimal structures were analyzed, these RNAs fell into different clusters, 3 of which represented most of the 6S RNAs. Information from these 3 clusters was used to identify 6S RNAs across genomes. Through this clustering method, several 6S RNAs were obtained in *Mycobacteria* and *Streptomyces* species, representative of actinobacteria.

We observed a similar discrepancy in an important family of RNAs known to be targets of the ANTAR RNA-binding protein. RNAs bound by ANTAR proteins are conserved in structure and are widespread among firmicutes and proteobacteria (Chai and Stewart 1998; Drew and Lowe 1989; Wilson et al. 1996b; Goldman et al. 1994; Ueki and Lovley 2010; Ramesh et al. 2012). In actinobacteria, however, despite the widespread presence of ANTAR protein domains (Pfam: PF03861), their target RNAs remained unidentified. Only recently, in a study focusing on *Mycobacteria*, these RNAs were identified using a genome-wide covariance search approach combined with clustering (Mehta et al. 2020). A search model (structure based sequence alignment) enriched in firmicute and proteobacterial RNAs showed very high sequence and structure similarity and as a consequence failed to predict RNAs in actinobacteria. When diverse RNAs from different firmicutes and proteobacteria were added to the search model, they separated into several clusters based on sequence and structure similarity. This clustering resulted in a search model that successfully identified RNAs in *Mycobacteria* by removing the bias imposed by highly similar or highly dissimilar RNAs. Notably, neither the firmicute (Ramesh et al. 2012) nor the mycobacterial search models (Mehta et al. 2020) were however not effective in finding ANTAR RNAs across the actinobacterial phylum.

Here, we identify the repertoire of ANTAR-target RNAs across actinobacteria. To identify these RNAs we first developed an actinobacteria-centric search model which when used to search against all actinobacterial genomes, successfully identified ANTAR-target RNAs. We find that the family of ANTAR-target RNAs is present across all actinobacteria and co-occurs with ANTAR proteins. There are only a few examples of bacteria where despite the presence of ANTAR proteins, we are unable to identify RNA targets. These RNAs resemble ‘cis’ regulatory RNAs in their genomic locations, typically residing in the untranslated region (UTR) or near the start of a coding region. COG analysis of the genes distal to ANTAR-target RNAs reveals that these RNAs are associated with transport and metabolism of small molecule metabolites, ranging from amino acids to metal ions to diverse sugar substrates. Additionally, ANTAR-target RNAs also appear linked to genes encoding transcription factors that are known to modulate the expression of several transporters. Our study underlines the presence of the ANTAR protein-RNA regulatory system in actinobacteria, and its importance in governing the uptake and metabolism of a variety of nutrients. This approach of scanning an existing RNA family for sequence diversity and using that to find homologs in distant phyla may be broadly applicable to other RNA families.

## RESULTS

### Identifying ANTAR-target RNAs across phylum Actinobacteria

Analysis of the previously reported ANTAR RNAs revealed that ∼400 ANTAR-target RNAs are known in firmicutes and proteobacteria (Ramesh et al. 2012), and they are conserved in secondary structure with dual stem loops separated by a linker (Fig 1A). Each stem possesses a hexanucleotide loop where the first and fourth positions are conserved in sequence as an adenine (A1) and guanine (G4) respectively (Fig 1A). More recently, in a study focusing on ANTAR RNAs in *Mycobacteria*, a covariance-based computational approach was used to search for ANTAR RNAs. Here it was shown that a focused search model (a set of RNAs aligned based on similar secondary structure and sequence) consisting of highly similar firmicute/proteobacterial RNAs was unable to predict RNAs in *Mycobacteria*. This is likely due to a divergence of mycobacterial ANTAR RNAs from their firmicute/proteobacterial homologs. Only when the search model was modified to include more diversity that expands the sequence space (partially focused search model), was the search capable of finding RNAs in *Mycobacteria*. This resulted in ∼90 ANTAR-target RNAs identified across all mycobacterial species (Mehta et al. 2020).

**Fig 1:**
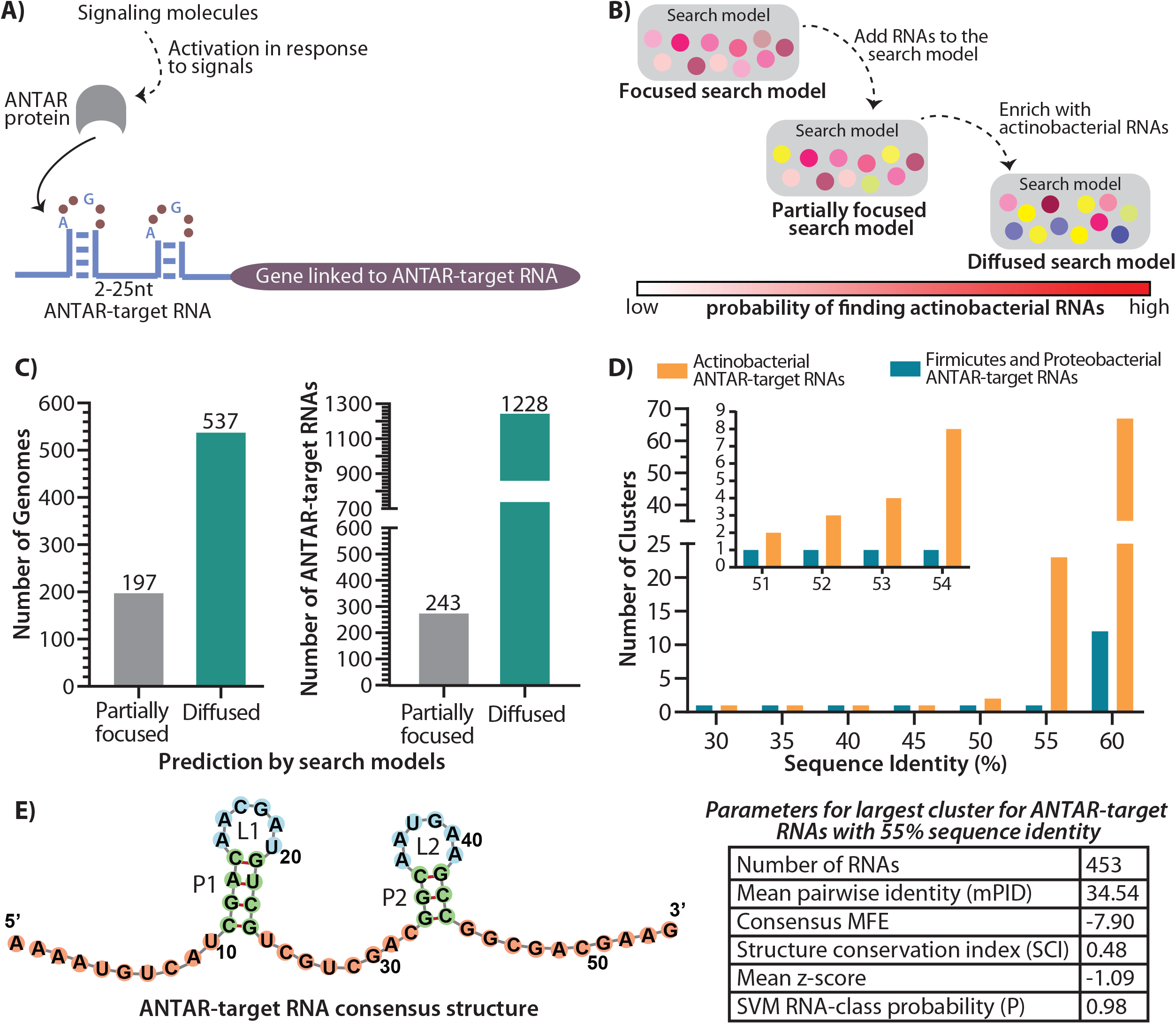
Improvised search model to predict ANTAR-target RNAs in actinobacteria. A) Cartoon showing the ANTAR protein-RNA regulatory system. Specific signals activate the ANTAR protein (grey), which upon activation binds the dual stem loop ANTAR-target RNA (blue). This results in regulation of the downstream gene (gene linked to ANTAR-target RNA, shown in purple). B) Schematic shows the steps performed to identify ANTAR-target RNAs using a covariance-based computational search. Previously reported search models with too little diversity (focused) did not yield any results in actinobacteria, while a search model with only moderate diversity (partially focused) identified ∼243 RNAs in actinobacteria, with a bit score threshold≥14. 30 actinobacterial representative RNAs from this set were used to enrich the search model further and this actionbacteria centric search model (diffused search model) resulted in a comprehensive list of ANTAR-target RNAs in actinobacteria. The probability of finding RNAs in actinobacteria is represented as a bar (red indicates high probability). C) Bar plot (left) shows the total number of actinobacterial genomes where RNAs are predicted using two different search models (gray and green). Bar plot (right) shows the total number of RNAs predicted using two different search models. The diffuse search model is able to predict RNAs in more than 60% of actinobacterial genomes as compared to the partially focused search model. D) RNA sets from firmicutes/ proteobacteria and actinobacteria were clustered using cmbuild. Bar plot shows the number of clusters obtained with varying sequence identity cut-offs imposed using cmbuild. Clusters obtained using 51%-54% sequence identity cut-off are shown as an inset. E) Consensus structure obtained for the actinobacterial ANTAR-target RNA sequences from the largest cluster with 55% sequence identity is visualized using Forna (Left). Stems are shown in green while the internal loops are shown in blue and the unpaired nucleotides are shown in pink. Parameters obtained from RNAz for the largest cluster with 55% sequence identity are shown (Right).

To further identify ANTAR-target RNAs in actinobacteria, we used the partially focused search model developed in the mycobacterial study (Fig 1B) and performed a covariance-based RNA search against all sequenced (∼720) actinobacterial genomes. This search could identify ∼243 ANTAR-target RNAs with high confidence. However, these newly found RNAs were restricted to less than 30% (197 of 720) sequenced actinobacterial genomes (Fig 1B-C).

In order to improve the search and predict RNAs more comprehensively across actinobacteria, we picked 30 representative RNAs from the initial 243 hits and created a new and fully actinobacterial search model (Table S1). This model was then used as input in a covariance search against the 720 genomes. This search now predicted ∼1288 RNAs with high confidence (Table S2 and S3), and importantly-RNAs were found in nearly 74.5% of sequenced actinobacteria (Fig 1B-C). Removal of identical RNAs from different strains of a species resulted in ∼611 unique ANTAR-target RNAs. Moreover, the 243 RNAs predicted initially, were also recovered in this search. This includes ANTAR-target RNAs predicted in mycobacterial species, which have been experimentally validated as binders of ANTAR proteins (Mehta et al. 2020).

We additionally analyzed this search model using the cmbuild program (Nawrocki et al. 2009), which creates a statistical profile of alignments and thus reports on the extent of sequence conservation and base-pairing potential (co-variation) within the aligned RNAs. Based on sequence (42% sequence identity) and structure (Covariance Model, CM score=0.48), the actinobacterial seed alignment shows significantly higher variation than the partially focused firmicute/proteobacteria seed (51% sequence identity and a CM score of 0.61). These results indicate that an actinobacteria-enriched search model that allows higher sequence/structural diversity while maintaining the core defining features of the RNA family is ideal for identifying new RNAs in actinobacteria.

In order to understand the characteristics of ANTAR-target RNAs in actinobacteria, we compared the 611 predicted actinobacterial RNAs with the previously reported 306 ANTAR-target RNAs from firmicutes and proteobacteria (Ramesh et al. 2012). Using cmbuild the RNAs from each set (actinobacterial versus firmicute-proteobacteria) were clustered at increasing sequence identity thresholds (Fig 1D). We find a stark difference between the two sets of RNAs. The actinobacterial RNAs start to separate out as clusters at a much lower sequence identity threshold (50%) when compared to firmicutes and proteobacteria (55%). This shows inherent diversity within the actinobacterial RNAs, possessing less than 50% sequence identity. We further analyzed the largest cluster of RNAs from each set for the extent of structural conservation. Even here, RNAs that are similar in sequence and hence clustered together showed a low CM (Covariance Model) score of ∼0.44 when compared to the firmicutes and proteobacterial set (CM score: ∼0.60). This confirms that actinobacterial RNAs allow for significantly higher sequence and structure variations (Fig S1A).

The largest cluster of actinobacterial RNAs represents ∼75% of the predicted RNAs. These were subjected to analysis using RNAz (Washietl et al. 2005; Altman et al. 2009) which computes a consensus secondary structure. We find that these RNAs, as expected fold into a dual stem-loop motif maintaining the core ANTAR-target RNA structural features, with a minimum free energy of −7.90 and a structure conservation index (SCI) of 0.48 (Fig 1E). The mean z-score of −1.09 obtained for these RNAs indicates that the structure motif observed is a stable true motif and does not occur by chance. The test for functionality based on SCI and z-score indicates that these RNAs belong to ‘functional RNA’ class (P>0.5). The consensus secondary structure for these RNAs shows more than 50% conservation of adenine and guanine in loop positions 1 and 4 respectively, and ∼50% conservation within the stems (Fig S1B).

### Distribution of ANTAR proteins and target-RNAs in actinobacteria

With a comprehensive list of ∼611 ANTAR-target RNAs identified, we looked at their distribution in the 128 known genera of actinobacteria and found that RNAs were predicted in genomes representing 87 genera which include 219 species (Fig 2A, inset). The majority of actinobacterial species possess 1 to 3 RNAs per genome (Fig 2A), while some species of *Actinomyces, Microbacterium, Bifidobacterium, Trupurella and Arthrobacter* appear to possess nearly 10 or even up to 36 different RNAs in the same genome (Fig 2A-B).

**Fig 2:**
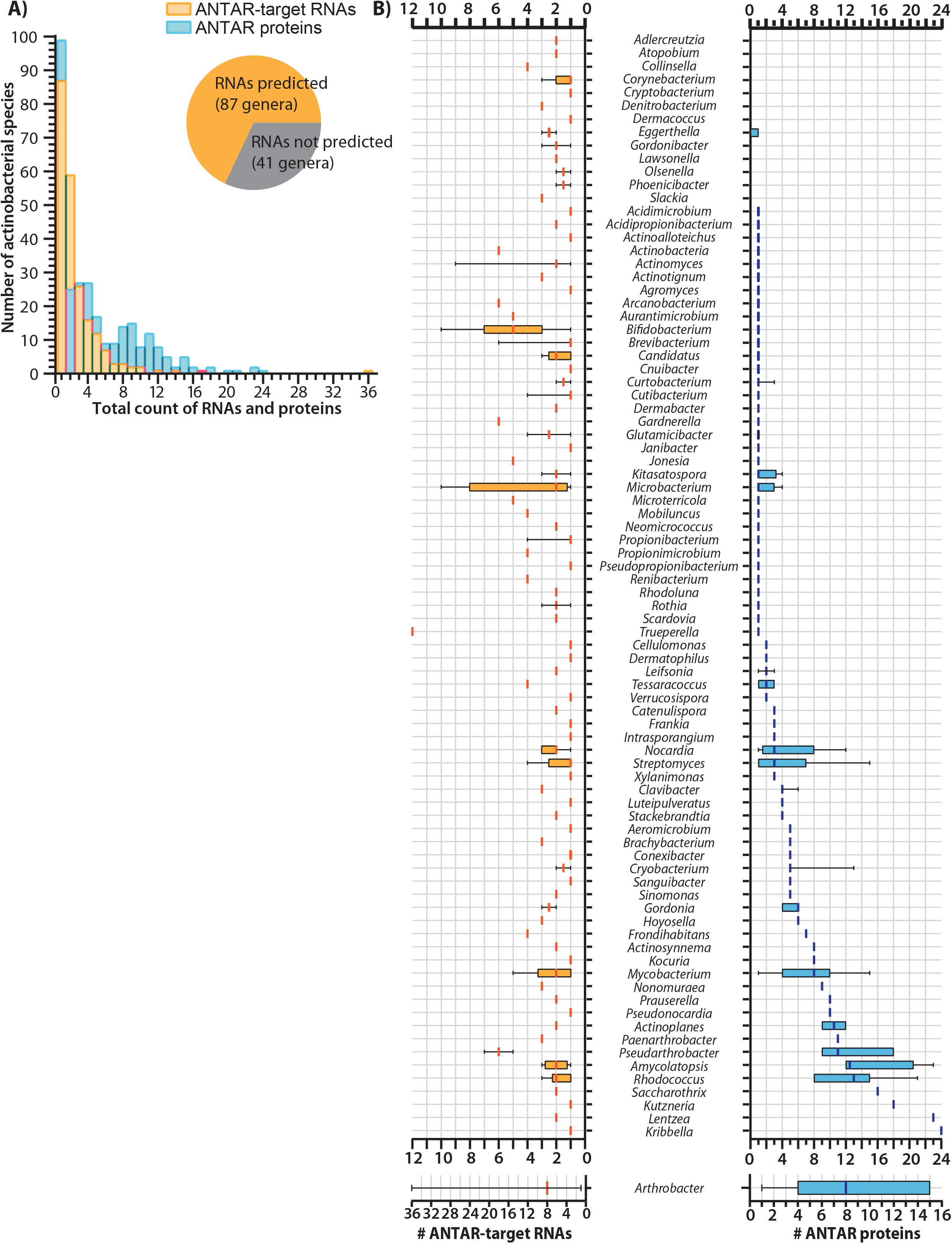
Distribution of ANTAR-target RNAs identified in Actinobacteria. A) Distribution of ANTAR-target RNAs (yellow) or ANTAR proteins (blue) in actinobacterial species. Inset pie-chart shows the number of actinobacterial genera where ANTAR-target RNAs are predicted. B) Distribution of ANTAR-target RNAs (left) and ANTAR proteins (right) in actinobacterial genera are shown as box-whisker plots. Median (vertical line), interquartile range (box) and 1.5 times the inter-quartile range (whiskers) are shown.

The ANTAR domain is an RNA-binding domain and proteins containing this domain are known to selectively recognize and bind RNAs of this family. Hence we asked if the distribution of RNAs reflected the distribution of the ANTAR proteins. To this end, we performed an HMMsearch using the ANTAR domain HMM model from the protein family database (Pfam: PF03861). With an e-value threshold set to 1e-4, we identified ∼1459 ANTAR-domain containing proteins in 245 species of actinobacteria. As seen for the RNAs, the distribution of ANTAR proteins too shows high variation, ranging from 1 to greater than 10 ANTAR proteins in a genome (Fig 2B, Fig S2). Interestingly, within the same genome, we do not always see a one to one correlation between the number of RNAs predicted and the number of ANTAR proteins present (Fig 2B). For example, in *Xylanimonas* there are 3 distinct ANTAR domain proteins with unique domain architectures. However, we predict only one ANTAR target RNA here, suggesting that the same RNA may act as a hub through which many different ANTAR proteins may act, towards different cellular outcomes. In contrast, *Trueperella* appears to possess a single ANTAR domain protein but 12 predicted RNAs, suggesting that many convergent processes may be controlled by ANTAR in *Trueperella*.

We found examples (<30% of species) where no RNAs were predicted despite the presence of one or more ANTAR proteins in the genome. Similarly, in a few examples no ANTAR proteins are present in a genome even though ANTAR-target RNAs are predicted with high confidence. Whether or not ANTAR proteins and RNAs have an active role in these organisms, or if alternate approaches are required to find RNAs and proteins in these organisms remains to be seen (Fig 2B, Fig S2). Regardless, these analyses imply that within phylum actinobacteria there is diversity of ANTAR function and mechanism.

### ANTAR-target RNAs are located in untranslated and coding regions of mRNAs

Previous studies have shown that ANTAR proteins, upon activation (through phosphorylation) bind to their target-RNAs and regulate downstream gene expression in *cis (Ramesh et al. 2012; Fox et al. 2009; Wilson et al. 1996a; Chai and Stewart 1998; Mehta et al. 2020; Malaka De Silva et al. 2020; Weber et al. 2019)*. Hence we analyzed the genomic locations and contexts of the predicted RNAs.

Based on genomic location, RNAs were categorized as: 1) intergenic (RNA lies 15nt-500nt upstream to an ORF), 2) sequester RBS or AUG (RNA harbors the ribosome-binding site (RBS) or the start codon or 3) inside ORF (RNA resides after the ORF start-site and lies within ≤100nt of the ORF start-site) (Fig 3A-B).

**Fig 3:**
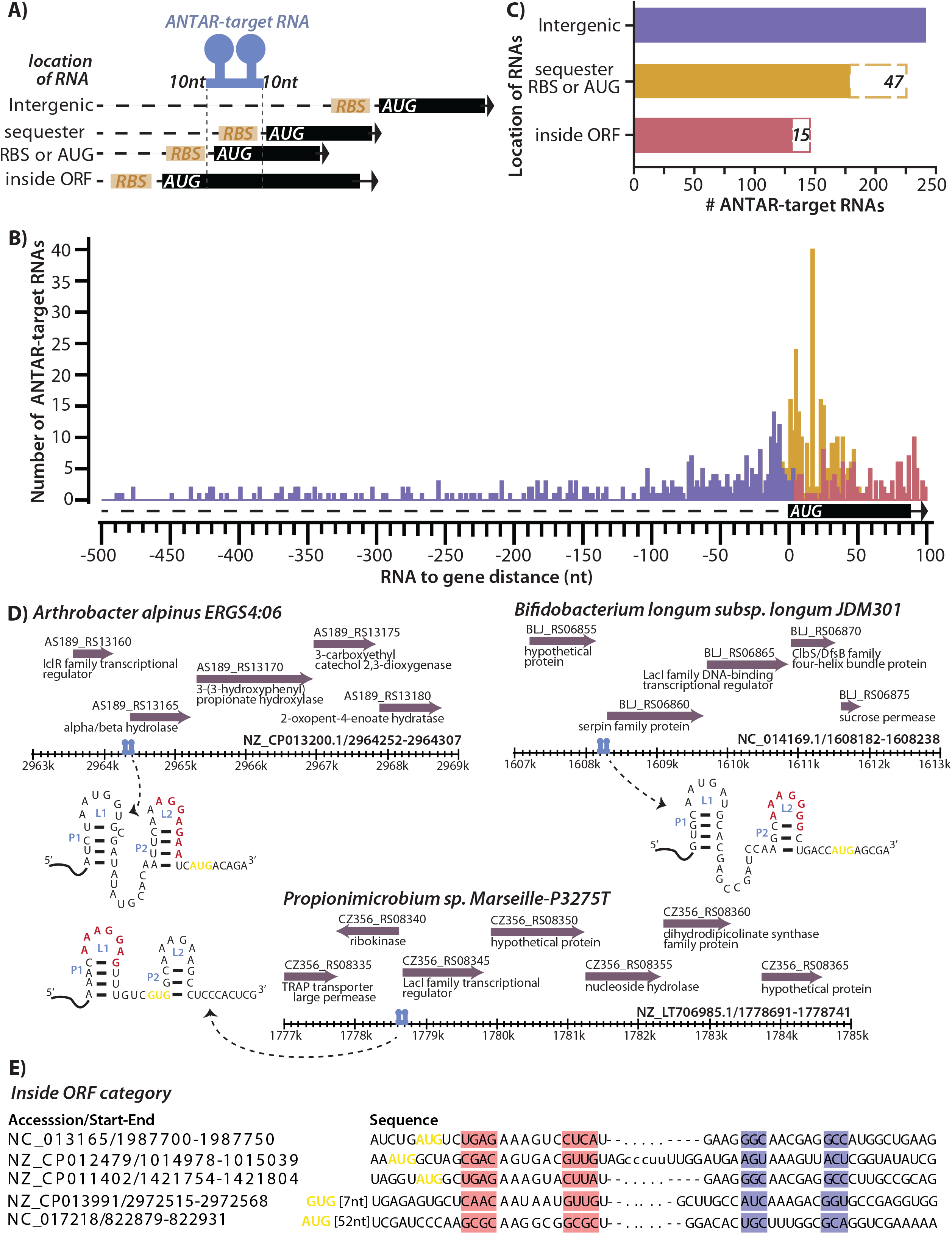
Locations of ANTAR-target RNAs within their genomic context. A) Schematic shows the location of ANTAR-target RNAs. RNAs are grouped in three categories-intergenic’ for RNAs that lie at a distance>15nt from start of ORF, ‘sequester RBS or AUG’ for RNAs which overlap with the ribosome binding site or start codon and ‘inside ORF’ for RNAs which lie after the start codon. 10-nucleotide flanking regions on either side of the dual stem loop structure are included in the distance calculations. B) Histogram shows distribution of RNAs versus their distance from the respective ORF. Several RNAs are found near the ORF start site, sequestering either RBS or AUG (yellow). C) Plot shows total number of predicted RNAs in three categories as described in panel A. 47 RNAs (dashed brown box) in the ‘sequester RBS or AUG’ category and 15 RNAs (dashed red box) in the ‘inside ORF category were assigned based on alternate ORF predictions. D) Representative RNAs from ‘sequester RBS or AUG’ category are shown with the ANTAR-target RNA structure marked. Potential RBS (red) and start codon (yellow) are shown. Genomic context of these RNAs (blue) are shown with ORFs (purple) with their NCBI gene annotations. E) Representative RNAs from the ‘inside ORF’ category are shown. The dual stems of the ANTAR-target RNA are highlighted in pink and blue. Start codon is marked in yellow.

We find that from a total of 611 RNAs analyzed, ∼39% RNAs are intergenic with a majority lying immediately upstream of an ORF, possibly in the 5’UTR of the corresponding mRNA (Fig 3C, Table S4). These RNAs were subjected to rho-independent terminator prediction using TransTermHP v2.08 (Kingsford et al. 2007) but only few of the RNAs appear to reside upstream of a terminator, with the second stem loop showing alternate base-pairing with the terminator (Fig S3). These few examples are reminiscent of ANTAR-target RNAs in firmicutes and proteobacteria, where binding by the ANTAR protein stabilizes the two-stem loop anti-terminator structure, allowing transcription of the downstream gene. With high GC genomes, it is possible that terminator predictions are inaccurate for these bacteria and hence other approaches may be required to ascertain the mode of transcriptional regulation.

Nearly ∼37% of the actinobacterial target-RNAs overlaps directly with the RBS or start codon (Fig 3C, Table S4). In a recent report it was shown that in *M. tuberculosis* and *M. smegmatis*, binding of activated ANTAR protein to such target RNAs, represses translation of the downstream mRNA, possibly by occluding the ribosome from binding the RBS (Mehta et al. 2020). We see similar features in these RNAs. For example, in *Arthrobacter alpinus* and *Bifidobacterium longum*, the RBS is sequestered within the second stem-loop, whereas in *Propionimicrobium species* both the RBS and the ORF start site lie within the ANTAR-target RNA (Fig 3D).

The ‘inside ORF’ category consists of ∼24% ANTAR-target RNAs (Fig 3C, representatives shown in Fig 3E). Several studies on non-coding RNAs (Tapsin et al. 2018; Del Campo et al. 2015; Tsuchihashi and Kornberg 1990; Chen et al. 2013; Gorochowski et al. 2015; Bandyopadhyay et al. 2020; Murat et al. 2014; Caliskan et al. 2015; Giedroc and Cornish 2009) have shown that structured motifs within the mRNA transcript may influence mRNA stability or regulate translation. It is possible that these ANTAR-target RNAs also control gene expression, though the detailed mechanism needs to be uncovered.

### Cellular pathways and genes associated with actinobacterial ANTAR-target RNAs

We next asked what cellular processes are linked to ANTAR in actinobacteria. Studies in *Enterococcus, Pseudomonas, Klebsiella, Acinetobacter* and *Geobacter* reveal that ANTAR-target RNAs are linked to nitrogen utilization (Ramesh et al. 2012; Drew and Lowe 1989; Chai and Stewart 1998; Malaka De Silva et al. 2020). Only few studies in actinobacteria have investigated the role of ANTAR. In *Mycobacteria*, ANTAR mediated gene regulation might influence lipid and related redox processes (Mehta et al. 2020) while a recent study in *Streptomyces*, show that the deletion of ANTAR-protein (*SSDG_04087*) impairs the developmental process and antibiotic production (Li et al. 2020). For this analysis, we took genes upstream of, or harboring ANTAR-target RNAs as input and performed COG analyses using the eggNOGmapper server. eggNOGmapper is a tool that performs a protein sequence homology search against precomputed eggNOG protein database to identify orthologs using a BLAST-like approach, and assigns the COG functional categories, KEGG pathways and gene ontology terms from orthologs to the query (Huerta-Cepas et al. 2019, 2017).

Our analysis showed that ∼85% of ANTAR linked genes belong to 17 different COG categories, while 15% are genes of yet unknown function (Fig 4A-B, Table S4). The majority of genes encode proteins involved in transport and metabolism of compounds, with a smaller subset restricted to enzymes involved in energy production. Core cellular processes including transcription, translation, replication and DNA repair also appear to be linked to ANTAR-target RNAs, and make up the next largest categories of COGs (Fig 4A). Additionally, we find a diversity of metabolites whose transport and metabolism would be linked to ANTAR (Fig 4B), with carbohydrate, amino-acid and lipids standing out as preferred metabolites.

**Fig 4:**
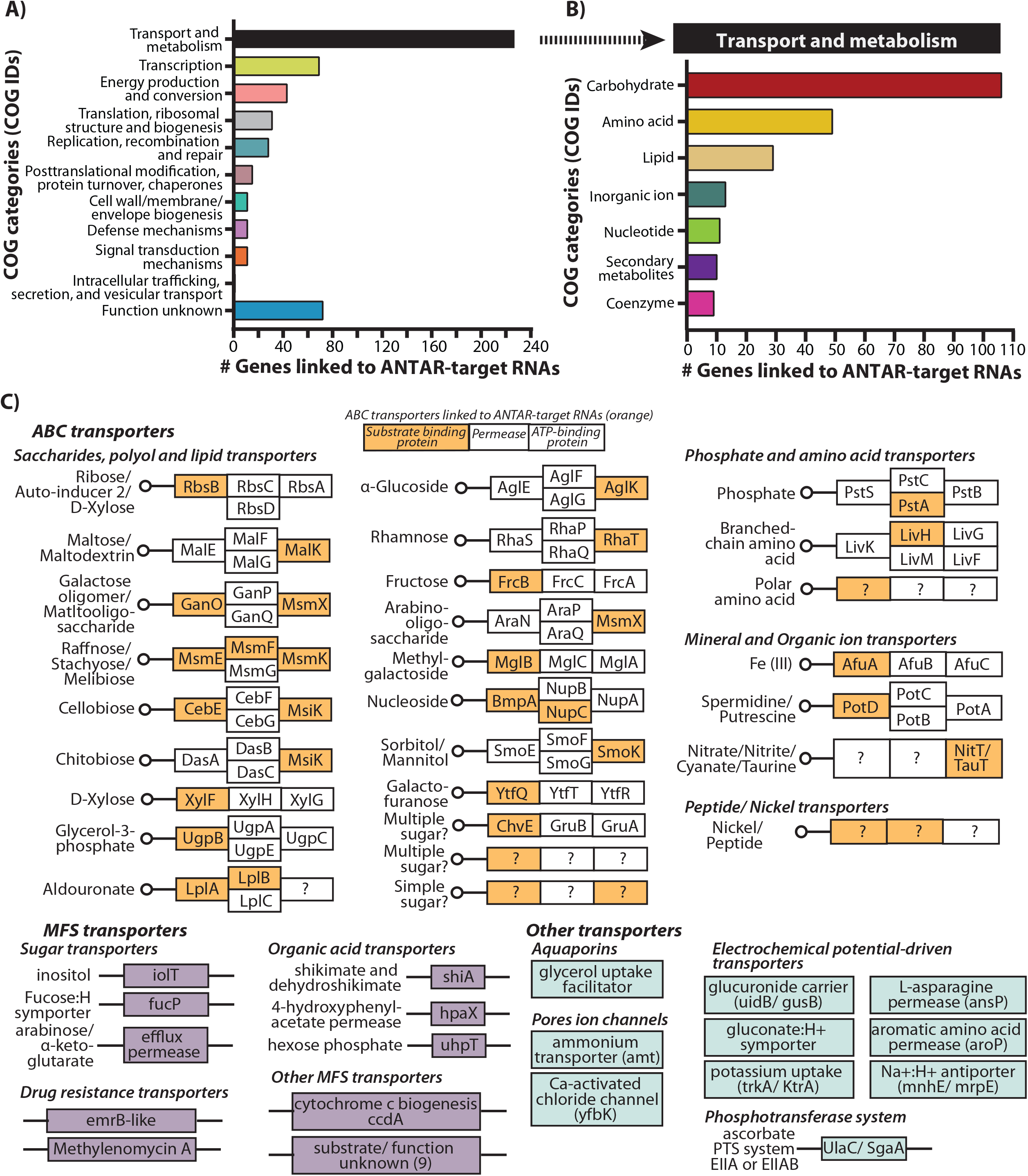
COG analysis of genes linked to ANTAR-target RNAs in actinobacteria. A) Genes linked to ANTAR-target RNAs, analysed using EggNOG-mapper, get assigned to 11 COG categories. Bar plot shows distribution of genes linked to ANTAR-target RNAs, in each COG category. B) Bar plot shows distribution of genes linked to ANTAR-target RNAs, within the ‘transport and metabolism’ COG category. Carbohydrate and amino-acid transport and metabolism are the major processes represented by the targets. C) ABC transporters with the substrate binding protein, membrane bound permease and ATP-binding components (boxes) are shown. Components of the transporter whose functions are not known are marked (?). Transporter components whose transcripts harbor an ANTAR-target RNA are marked in orange. Genes linked to ANTAR-target RNAs, encoding MFS transporters (purple) and other transporters (blue) are shown.

We next asked if a cellular process or function linked to ANTAR was restricted to any particular branch within the actinobacterial phylogenetic tree (Fig S4). Some processes such as replication, recombination and repair and transcription, are ubiquitously seen linked to ANTAR, in most genera. In contrast, intracellular trafficking, secretion and vesicular transport process appear restricted to *Gordonia* species while translation related processes and lipid transport and metabolism are largely restricted to non-pathogenic *Mycobacterium* and *Nocardia* species respectively. Energy production and conversion is found to be conserved in species of *Pseudarthrobacter, Renibacterium, Sinomonas, Rhodococcus, Gordonia and Mycobacterium*.

We next asked if closely related genera have co-opted ANTAR for similar processes. Indeed, several species from *Bifidobacterium* and *Gardnerella*, have processes such as carbohydrate transport and metabolism, transcription, translation related processes and cell-membrane biogenesis linked to ANTAR. Similarly, six processes including cell energy production and conversion process, transcription and translation related processes, signal transduction mechanisms and amino-acid and carbohydrate transport and metabolism are linked to ANTAR in closely related *Arthrobacter* and *Pseudoarthrobacter* species (Fig S4).

KEGG pathway and KEGG BRITE analysis of transporters whose genes are linked to ANTAR, show that they belong to the ABC transporter, MFS sugar transporter and Aquaporin families (Table S5). The ABC transporter complex consists of multiple components: a periplasmic substrate-binding protein, one or more trans-membrane permeases, an ATP-binding protein and occasionally a substrate-specific enzyme (Higgins 1992). Interestingly, we find that different components of the transporters, especially the substrate recognizing proteins harbor the ANTAR-target RNA in their mRNA (Fig 4C). This makes intuitive sense since transporters are often under tight regulation and the different components are made only upon sensing the presence of the cognate sugar/metabolite.

The second highest COG category is that of transcription with over 19 different transcription factor families linked to ANTAR-target RNAs (Fig S5A). Remarkably, the majority of these transcription factors are known to regulate the expression of transporter proteins, once again tying back ANTAR-target RNAs to the transport of small molecule metabolites. ∼29% of transcription factor encoding genes linked to an ANTAR-target RNA belong to the LacI type transcription factor that are major regulators of sugar catabolic genes (Fig S5A). For example, in *Streptomyces lydicus*, a LacI transcription factor (TF) carrying an ANTAR-target RNA is present upstream of transporter components involved in ribose uptake (Fig S5B). In *Corynebacterium glutamicum*, the homologous TF (*cg1410*) is reported to regulate the downstream *rbsDACBK* operon in response to ribose availability (Nentwich et al. 2009). Similarly, even in *Bifidobacterium dentium, Gardnerella vaginalis* and *Microbacterium sp*. and *Cryobacterium arcticum*, ANTAR-target RNAs are also linked to LacI transcription factors that regulate other sugar transporters and sugar related genes (Table S2 and Table S4). These results suggest that in several actinobacteria sugar transporters, as well as the proteins regulating sugar transport and metabolism are under the influence of ANTAR regulation.

TetR family TFs are also linked to ANTAR-target RNAs in several actinobacterial species (Fig S5A). Transcription factors belonging to this family typically regulate the expression of enzymes from different catabolic pathways or proteins involved in multi-drug resistance (Table S2 and Table S4) (Ramos et al. 2005; Bhukya and Anand 2017). An ANTAR-target RNA in *Mycobacterium marinum* is found upstream to the *MMAR_RS11360* gene encoding a TetR family transcription factor (Fig S5B). Its *M. tuberculosis* homolog, *Rv1474c* is found to cotranscribe with the upstream aconitase gene and regulates aconitase expression in response to iron (Balakrishnan et al. 2017). A conserved operon in *Streptomyces* species is predicted with an ANTAR-target RNA upstream to a *SufR* encoding gene, *SACTE_RS06635* (Fig S5A-B). *SufR* is an ArsR family transcription factor and a repressor of the downstream *sufBCDS* operon, the primary Fe-S assembly cluster system, that responds to the availability of Fe-S cluster required as protein cofactors in many cellular processes (Cheng et al. 2020). Both these examples underline that an additional layer of post-transcription gene regulation is likely imposed by virtue of ANTAR-target RNAs.

## DISCUSSION

In this study, we have developed an actinobacterial centric computational search model that helped us identify the repertoire of ANTAR-target RNAs in phylum actinobacteria. These RNAs are distinct from their firmicute and proteobacterial homologs in sequence, while still maintaining the conserved dual stem structure so characteristic of ANTAR-target RNAs. Interestingly, several of the actinobacterial RNAs overlap with the ribosome binding site or the start codon of the downstream mRNA, suggesting that translational control may be a prominent mode of regulation by ANTAR-target RNAs. Through extensive analysis we show that the genes linked to ANTAR RNAs are likely to regulate the transport and metabolism of small molecule metabolites, providing insights into the cellular pathways they may influence.

Key to our findings was the development of a novel computational search model that was effective in identifying these structured dual stem loop RNA motifs. Covariance search programs rely on both the sequence and the base-pairing information within a search model, to find similar RNA motifs in a genome. Previously reported ANTAR RNA search models (Ramesh et al. 2012; Mehta et al. 2020) either failed or were only partially successful in predicting RNAs in actinobacterial genomes due to a lack of diversity in sequence and base-pairing potential. Removing the bias from highly similar or dissimilar sequences, the new search model developed in this study shows more sequence and structure diversity as compared to the previous models and this was key in identifying ANTAR-target RNA motifs in actinobacteria.

Analysis of the genomic locations of ANTAR-target RNAs from actinobacteria reveals many examples where the RNA is next to the ORF start site, either sequestering the RBS or start codon within the dual stem motif of the ANTAR RNA. A similar genomic arrangement of ANTAR RNAs was seen previously in *Mycobacteria (Mehta et al. 2020)*, was shown to function via translational repression. Here, RNAs bound by activated ANTAR protein were shown to repress translation, possibly by preventing ribosomes from accessing the RBS. Our analysis indicates that translational control via ANTAR-target RNAs may be a prominent mode of regulation in actinobacteria.

Analysis of cellular processes likely to be controlled by ANTAR-target RNAs revealed a link between these RNAs and the transport and metabolism of small molecule compounds, especially carbohydrates, amino acids and lipids. Certain species of *Bifidiobacterium, Gardnerella* and *Scardovia* show conservation of ANTAR-target RNAs in transcripts encoding carbohydrate transport and metabolism proteins. For these genera, sugar utilization is intricately linked to physiology. For example, *Bifidobacteria* are sacchrolytic intestinal bacteria detected in human and animals (Pokusaeva et al. 2011), while *Scardovia* is detected in human dental caries and adeptly use carbohydrate fermentation pathways to lower the pH of the oral biofilm and likely induce caries progression in the host (Kressirer et al. 2017; Kameda et al. 2020). Pathogenic *Gardnerella vaginalis* have the ability to degrade glycans in the host mucosal epithelial layers to invade and colonize in the host (Lewis et al. 2013). Species belonging to genus *Nocardia* shows that ANTAR-target RNAs might regulate lipid transport and metabolism similar to to that seen in *Mycobacteria*. Our results link ANTAR-target RNAs to metabolite transport and utilization in these organisms, possibly indicating that ANTAR regulation may contribute to their growth and survival within their host.

An important finding from our study is the association of ANTAR-target RNAs with mRNAs encoding transcription factors. Transcription factors themselves are regulators of gene-expression, often regulating multiple target genes. By controlling the expression of a transcription factor, even a single ANTAR-target RNA in the genome could indirectly control the expression of multiple genes. We also observed that many of the transcription factors whose mRNAs harbor ANTAR-target RNAs, in fact regulate sugar and other metabolite transport. This implies that the scope of ANTAR-based control of metabolite transport is much broader. ANTAR targets are also involved in metabolic pathways that are wide spread in the utilization of sugar as an energy source.

Additionally ANTAR is involved in the production of essential amino acids, nucleotides or involved in biosynthesis of antibiotics. Actinobacteria thus seem to have co-opted ANTAR-based regulation for diverse metabolic pathways according to their growth requirements in the diverse niches.

In a recent study in *Streptomyces pristinaespiralis*, deletion of the ANTAR protein *SSDG_04087* led to a bald phenotype (loss of hyphae formation) and reduced production of the antibiotic pristinamycin (Li et al. 2020). In our study, we identify four ANTAR-target RNAs in *S. pristinaespiralis*, one of which lies in the transcript of a sugar (fructose) transporter protein (*SPRI_RS32325*). The uptake of complex sugars by *Streptomyces* favors development (sporulation) and production of antibiotics (Rueda et al. 2001; Swiatek et al. 2012; Sciences and 2013). In fact, perturbation of glycolysis/ gluconeogenesis pathways is a standard method by which to increase the production of antibiotics by *Streptomyces*, for industrial applications (Butler et al. 2002; Li and Townsend 2006; Ryu et al. 2006; Huang et al. 2012). Another ANTAR-target RNA is found in the mRNA for the enzyme agmatinase (*SPRI_RS23705*), that converts arginine to putrescine. Putrescine is a precursor of succinate (Krysenko et al. 2017; Schneider and Reitzer 2012) that can feed into the TCA cycle and the synthesis of various amino acids, which are directly involved in the production of the antibiotic pristinamycin (Voelker and Altaba 2001; Zhang et al. 2012). The discovery of these ANTAR-target RNAs in *Streptomyces* thus implicates gene *SPRI_RS32325* and *SPRI_RS23705* as possible candidates that might be investigated to understand the observed phenotype. Our comprehensive description of ANTAR-target RNAs and ANTAR proteins in actinobacteria now provides a resource for microbiologists to mine.

## MATERIALS AND METHODS

### Actinobacterial genomes used in this study

720 actinobacterial genomes, with their corresponding gene annotations and proteomes are listed as “Complete genomes” in NCBI (RefSeq v92). These were considered in this study. Corresponding taxon IDs for these organisms were taken from NCBI and a taxonomy tree was retrieved in Phylip format from NCBI Batch Entrez (https://www.ncbi.nlm.nih.gov/sites/batchentrez). The phylogenetic tree visualization was carried out using iTOL (Letunic and Bork 2016).

### Predicting ANTAR-target RNAs in actinobacteria using covariance

A search model previously reported for identifying ANTAR-target RNAs in *Mycobacteria* (partially focused search model) (Mehta et al. 2020), was taken and an initial covariance search with a bit score threshold of 10.0 was carried out against actinobacterial genomes using Infernal v1.0.2 (Nawrocki et al. 2009). High confidence RNAs with a bit score≥14.0 and showing a dual stem loop structure (with at least 3 base-pairs in each stem and hexanucleotide loops allowing a single point variation) were considered as putative ANTAR-target RNAs. 30 of these predicted RNAs from actinobacteria were taken to form an actinobacteria centric search model (diffused search model). cmbuild analysis of the partially diffused and diffused search models reports on the CM (Covariance model) score where a higher CM score was taken as an indication of highly similar sequences. To identify ANTAR-target RNAs in actinobacteria, 720 genomes representing 315 actinobacterial species were subjected to covariance search using the diffused search model. Hits with a bit score threshold≥15 and lying between 500nt upstream to 100nt downstream of the nearest ORF were retained. RNAs that are identical to the search model or are single-nucleotide point variants were considered for further analyses. Redundant identical RNAs from strains were removed and unique RNAs from each species were considered as a representative. We used cmbuild (Nawrocki et al. 2009) and RNAz (Washietl et al. 2005; Altman et al. 2009) to analyze the predicted ANTAR-target RNAs for their sequence and structure similarity. Using the cmaxid option of cmbuild implemented in Infernal v1.0.2, we performed a clustering analysis. Sequence identity cut-off ranging from 30%-60% was imposed during clustering such that any two RNAs that have sequence identity more than the cut-off, willform a cluster reported with a corresponding CM score. Any group with <2 RNAs was not considered. The -cdump option of cmbuild writes the multiple sequence alignment for the clusters. The multiple sequence alignment of the largest cluster formed with 55% sequence identity cut-off, was further checked for functionality using RNAz. RNAz calculates i) the structure similarity of the individual RNAs to the consensus structure, reported as structure conservation index (SCI) ii) z-score that describes the standard deviation of the structures formed by the RNAs in a cluster against the structures for a random set of RNAs with same length and base composition, where the negative z-score indicates a true stable structure and has not occurred by chance. Based on these two measures, the RNAs with conserved and stable structures (P>0.5) are considered as a ‘functional RNA’ class. Consensus RNA structure for the largest cluster was visualized using forna (Kerpedjiev et al. 2015) and nucleotide-level resolution of the consensus structure was obtained using R2R (Weinberg and Breaker 2011) and statistically significant covarying positions were identified using R-scape (Rivas et al. 2016).

### Distribution of ANTAR domain containing proteins in actinobacteria

An HMM model for the ANTAR domain was taken from Pfam v33.0 (PF03861) and HMMsearch (hmmer v3.2.1) was performed against all actinobacterial proteomes with e-value threshold 1e-4. This identified proteins having ANTAR domains. Proteomes where the HMMsearch failed to identify ANTAR proteins, were further searched for sequences homologous to the Rv1626 ANTAR domain using BLASTp with evalue threshold 1e-3.

### Categorizing ANTAR-target RNAs based on location within the genomic context

ANTAR-target RNAs were grouped into 3 categories based on their distance from ORFs. RNAs (including 10nt flanking region) that are 15nt upstream from start of ORF, were assigned to ‘intergenic’ group. RNAs that completely reside within the ORF were assigned to ‘inside ORF’ group. RNAs that harbor a potential RBS as part of the RNA structure, are grouped as ‘sequester RBS or AUG’. RNAs were also subjected to alternate ORF (altORFs) prediction using standalone NCBI ORFfinder (https://www.ncbi.nlm.nih.gov/orffinder/) with default parameters allowing for ATG or any alternate start codons. Predicted ORFs which harbor a potential ribosome-binding site (RBS) with a 4-6nt AG-rich region and reside 0-15nt upstream of the start codon are considered as putative altORFs. RNAs from the ‘intergenic’ group were further subjected to Rho-independent terminator prediction. Here, target-RNA sequences along with 40nt downstream sequences were given to TransTermHP (Kingsford et al. 2007) with parameters uwin-require=0 and min-conf=50.

### COG and KEGG pathway analyses for ANTAR targets

Protein sequences of genes linked to ANTAR-target RNAs were subjected to COG analysis using EggNOG mapper v4.5.1 (http://eggnogdb.embl.de/#/app/emapper). A minimum 70% query coverage and e-value threshold 1e-3 was used to assign COG categories and KEGG orthologs (KO) based on sequence homology. Independently, these protein sequences were given as input to KofamKOALA (https://www.genome.jp/tools/kofamkoala/) with e-value threshold 1e-2, which reports on top KEGG orthologs using an HMMsearch. Orthologs for genes linked to ANTAR-target RNAs, were mapped using EggNOG and/or KofamKOALA (Table S4). These KOs were then given to KEGGmapper (“KEGG reconstruct pathway” and “KEGG search and color pathway”) for pathway analyses. Visualization of data was carried out in iTOL and the pathway graphs were obtained using KEGG and modified using Adobe Illustrator. All plots were obtained using Graphpad Prism v8.0.

## ACKNOWLEDGMENTS

We are grateful for funding and support from the DBT/Wellcome Trust-India Alliance (IA/I/14/2/501521), DST-SERB grant no. ECR/2016/001593 and the Human Frontier Science Program research grant RGY0077/2019 to A.R. We also acknowledge support from the Department of Atomic Energy, Government of India and the National Centre for Biological Sciences-TIFR, under project no. 12-R&D-TFR-5.04-0800. D.M. was supported by ICMR-SRF fellowship (No. ISRM/11(18)/2019. We are especially grateful to Dr. R. Sowdhamini (NCBS) and Dr. Sabarinathan Radhakrishnan (NCBS) for valuable suggestions on this work. We thank Anirudh KN and other members of the Ramesh Lab for feedback and suggestions.

